# Carbon isotope composition for agronomic diagnostic: predicting yield and yield response to nitrogen in wheat

**DOI:** 10.1101/2021.08.16.456583

**Authors:** C. Mariano Cossani, Victor O. Sadras

## Abstract

Rainfed crops rely on two sources of water: stored soil water at sowing and seasonal rain. In strongly seasonal winter-rainfall environments, stored soil water at sowing is minor, and uncertain seasonal rainfall is a source of risk. In south-eastern Australia, under-fertilisation is a common outcome of nitrogen risk management with implications for yield and soil mining. Here we explore the use of carbon isotope composition (δ ^13^C) to capture the effects of water in the prediction of yield and guide nitrogen management. In the sampled environment, crops receive at least 50% of seasonal rainfall by stem elongation, and at least 70% of seasonal rainfall by flowering. In a sample of 1518 plots, yield varied from 0.07 to 9.96 t ha^-1^ and correlated with δ ^13^C measured with isotope ratio mass spectrometer (IRMS) at flowering; this is consistent with the rainfall pattern and the physiology of the crop featuring a critical period for yield from 300 °Cd before to 100 °Cd after anthesis. In a sample of 135 plots, yield varied from 1.2 to 8.4 t ha^-1^ and correlated with δ ^13^C measured with IRMS at stem elongation. Yield response to nitrogen, defined as the difference between yield in fertilised crops (50 to 200 kg N ha^-1^) and unfertilised controls, correlated with δ ^13^C measured with IRMS at stem elongation, except for late-sown crops. Mid-infrared spectroscopy (MIR) returned estimates of δ ^13^C that agreed with δ ^13^C measured with IRMS (calibration: R^2^ = 0.82, RMSE = 0.53‰, n = 833; validation: R^2^ = 0.70, RMSE = 0.75‰, n = 364). We conclude that a MIR based, high-throughput, affordable measurement of δ ^13^C could be scaled to guide nitrogen management of wheat in winter-rainfall environments.

## 1. Introduction

Management of nitrogen is critical for economic and environmental outcomes in agriculture (Angus and Grace, 2017; Cassman et al., 2002; Clark and Tilman, 2017; Jobbágy and Sala, 2014; Sadras et al., 2016). With uncertain rainfall, yield is uncertain and nitrogen fertilisation is a risky investment in winter-rainfall environments of Europe (Basso et al., 2012) and Australia (Armstrong et al., 2019; Meier et al., 2021; Monjardino et al., 2015; Monjardino et al., 2013; Neuhaus and Sadras, 2018; Sadras et al., 2016). Nitrogen can be managed strategically, for example building soil nitrogen banks, and tactically (Basso et al., 2011; Meier et al., 2021). Tactically, rainfall forecasts have been advanced to guide nitrogen fertilisation with mixed results (Anwar et al., 2008; Asseng et al., 2012; Moeller et al., 2008). Direct measurement of crop water status to guide nitrogen application, the focus of this study, has received less attention. Pancorbo et al. (2021) measured nitrogen and water status of wheat with remote sensing, and highlighted the difficulty of unscrambling crop nitrogen and water status using only reflectance-based vegetation indices.

Carbon isotope composition (δ ^13^C) integrates the water status of plant tissue over time, and has been used to screen for water use efficiency in annual crops (Araus et al., 1998; Bort et al., 2013; Condon et al., 1993; Farquhar and Richards, 1984; Hall et al., 1994; Rebetzke et al., 2002; Richards et al., 1993). Carbon isotope composition has been used as an environmental marker in ecology (Kohn, 2010; Schulze et al., 2006a; Schulze et al., 2006b), but less so in agriculture. Standard methods to measure isotope composition are involved and costly (Kleinebecker et al., 2009), hence the need for high throughput, cost effective methods. Okano (1983) demonstrated the feasibility of measuring δ ^13^C with infrared absorption spectrometry in C3 and C4 species. Calibrations were reported that relate δ ^13^C with near infrared reflectance (NIR 4000–13,000 cm^-1^ or 750–2500 nm) spectroscopy in alfalfa and grasses (Clark et al., 1995) and in native plants of southern Chile (Kleinebecker et al., 2009). In grain crops, Ferrio (2007) used NIR to measure δ ^13^C of durum wheat grains, but the method has not been tested for vegetative tissues required for fertiliser decisions early in the season. Attenuated Total Reflectance Fourier Transform Infrared spectroscopy (ATR–FT-IR) and Mid Infra-Red (MIR, 4000–400 cm^−1^ or 2500–25000 nm) returned affordable, more rapid analysis (< 1 minute), with smaller and less involved samples, improved sensitivity, and the ability to analyse multiple physical states of many compounds compared to NIR (Bureau et al., 2019).

Here we explore the use of δ ^13^C to predict yield of rainfed wheat in a context of fertiliser management, and the feasibility of using MIR to estimate δ ^13^C. The rationale of our study has three elements. First, yield is a primary function of grain number; grain weight is secondary when the environment is the main source of variation of yield, but could be more important when genotypes drive variation, particularly in high yielding environments (Calderini et al., 2021; Slafer et al., 2014). Second, wheat grain number is responsive to stress in a developmental window from approx. 30 days before to 10 days after anthesis (Fischer, 1985) or 300 °Cd before to 100 °Cd after anthesis (Dreccer et al., 2007; Telfer et al., 2018). Third, the most severe drought in winter-rainfall environments of Australia has an onset at about 400 °Cd before flowering and thus overlaps with the critical period, reducing grain number and yield (Chenu et al., 2013). From these three observations, we hypothesise that grain yield correlates with δ ^13^C in the critical stages under the prevailing agronomic conditions of winter-rainfall environments of south-eastern Australia, and that δ ^13^C could be used to inform nitrogen fertilisation.

## 2. Method

### 2.1. Experiments

Table 1 summarises the experiments used to source yield data and samples of plant material for δ ^13^C. Field trials were conducted in South Australia and sources of variation included 20 photothermal conditions (locations, seasons) combined with cultivars, initial soil nitrogen, fertiliser rate, initial soil water and growing season rainfall. Crops were grown on Dermosol and Calcarosol soils at Hart (Lat: -33.758, Long: 138.415), Sodosol and Chromosol at Turretfield (Lat: -34.553, Long: 138.786), Calcarosol at Roseworthy (Lat:-34.544, Long: 138.688) and a Vertosol at Mintaro (Lat: -33.884, Long: 138.774). The experiments were designed with specific objectives as described elsewhere (Cossani and Sadras, 2021; Hoogmoed et al., 2018; Hoogmoed and Sadras, 2018), and sampling for δ ^13^C was opportunistic (Table 1). We used plot as experimental unit for paired measurements of yield and δ ^13^C to capture both the original sources of variation (Table 1) and the variation between plots.

**Table 1.**
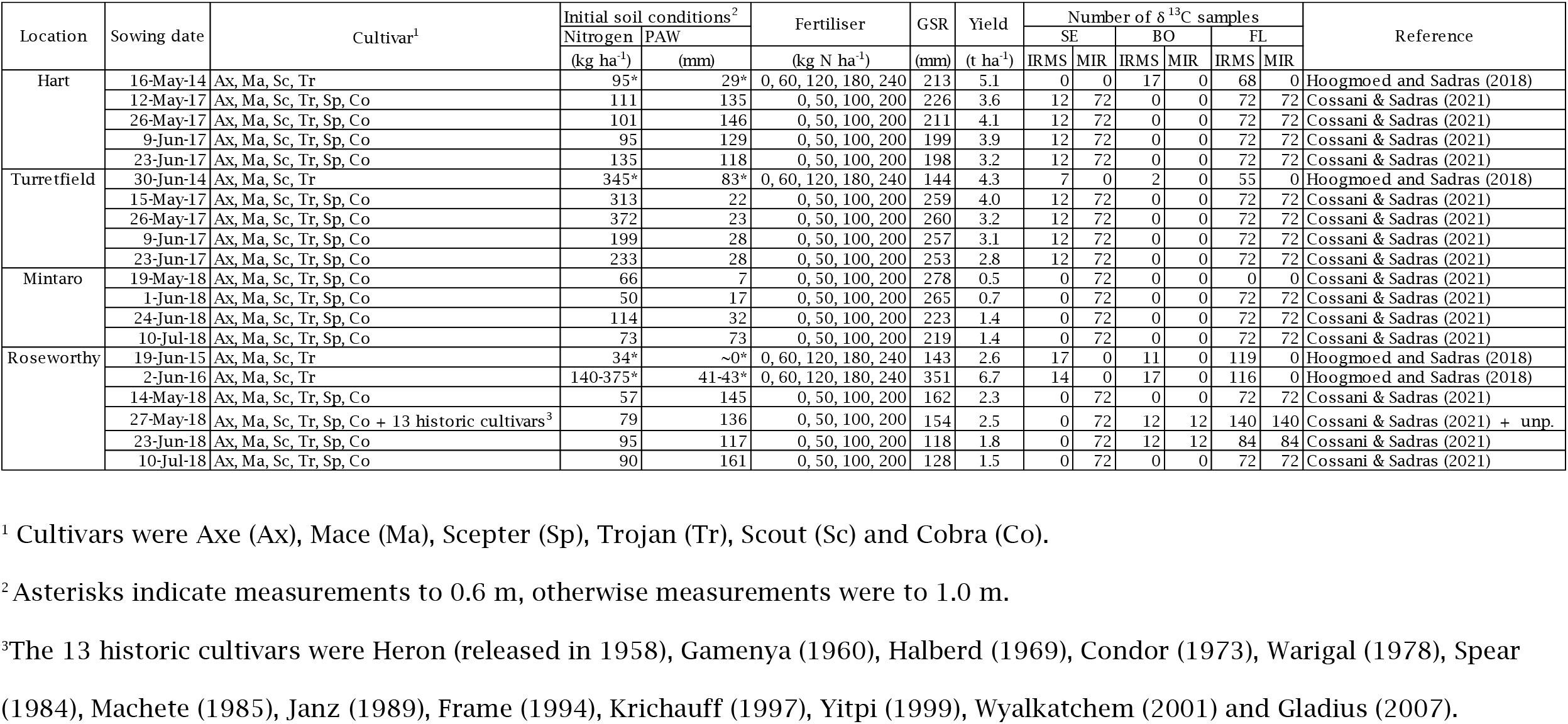
Locations, sowing dates, cultivars, initial soil nitrogen and plant available water (PAW), fertiliser rate, growing season rainfall (GSR) and range of yield in field experiments used to sample for δ ^13^C at stem elongation (SE, DC 2.3 to 3.2), Booting (BO, DC 3.7-4.9) and flowering (FL, DC 6.0 to 6.8). IRMS is isotope ratio mass spectrometer and MIR is mid-infrared spectrometry.

Crop phenology was monitored using the scale of Zadoks et al (1974). At beginning of stem elongation, booting, flowering and maturity, we sampled shoots in centre rows (2 × 0.5 m sections). Shoots were oven-dried at 65 °C for 48 hours and grinded with a mill (Thomas Wiley^®^ model 4, Swedesboro, NJ, United States) and a mesh for keeping particles smaller than 1 mm.

## 2.2 Carbon isotope composition

We measured δ^13^C in three steps. First, a total of 1724 samples (8% stem elongation, 4% booting, 88% flowering) was measured with an Isotope Ratio Mass Spectrometer (IRMS, Thermo Electron, Bremen, Germany) to probe for associations with grain yield (Table 1). Second, we scanned a total of 2408 samples with MIR spectroscopy using an ATR–FTIR spectrometer (Bruker ALPHA II, city, country); samples were scanned at constant temperature at a resolution of 4 cm^−1^ in the wavenumber between 4000 and 400 cm^−1^ with 24 scans per sample with air as background. Third, a subset of 1197 samples (92% flowering, 1% booting, and 7% stem elongation) from Step 2 was used to compare IRMS and MIR readings. Calibration and test spectral data were pre-processed using first derivative + vector normalization (SNV) and 17-point smoothing and mean centring. To perform the test set validation, 833 spectra were selected as calibration spectra and 364 different spectra were selected as test spectra. The calibration model was built using PLSR in the QUANT package of OPUS Version 7.0.129. The mean prediction error of the calibration was estimated using the RMSEE (root mean square error of estimation) calculated from the SSE (sum of the square errors), with M number of spectra and R number of PLS vectors namely rank:

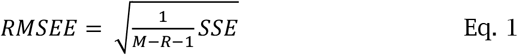

for the validation, we calculated the Root Mean Square Error of Prediction RMSEP:

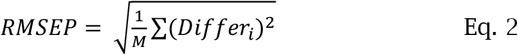

where *Differi* is the difference between the δ13C of sample *i* measured with IRMS and its δ13C predicted value. For both calibration and validation, the RPD (Residual Prediction Deviation) was estimated as the ratio of standard deviation to standard error of prediction.

### 2.3. Associations between yield and yield response to nitrogen with δ^13^C

We used regression analysis to investigate the associations of yield and yield response to nitrogen with δ^13^C. Yield response to nitrogen was calculated as the difference in yield between fertilised plots (50 to 200 kg N ha^-1^) and unfertilised controls. We fitted Reduced Maximum Axis (Model II) regressions necessary to account for error in both *x* and *y* (Ludbrook, 2012). We report *p*-value as a continuous quantity, and Shannon information transform [*s* = -log2(*p*)] as a measure of the information against the tested hypothesis (Greenland, 2019). Although *s* is a function of *p*, the additional information provided is not redundant. With base-2 log, the units for measuring this information are bits (binary digits). For example, the chance of seeing all heads in 4 tosses of a fair coin is 1/2^4^ = 0.0625. Thus, *p* = 0.05 conveys only *s* = −log2(0.05) = 4.3 bits of information, “which is hardly more surprising than seeing all heads in 4 fair tosses” (Greenland, 2019).

## 3. Results

### 3.1. Rainfall patterns: crops received at least half of seasonal rainfall by stem elongation and at least 70% by flowering

Climatically and agronomically, a rainfall season is defined for our target region that spans from April to October (French and Schultz, 1984; Gentilli, 1971). Long-term (1960-2020) seasonal rainfall averaged from 326 mm at Roseworthy to 448 mm at Mintaro. Actual seasonal rainfall sampled in the experiments varied ∼3-fold, from 118 to 351 mm (Table 1). Intra-seasonal variation was similar among locations: April rain contributed 0.10 of seasonal rainfall, which peaked at 0.17 midseason and dropped to 0.10 in October (inset Figure 1A). This intra-seasonal variation was reflected in curves relating cumulative fraction of seasonal rain and fraction of season crossing over the *y = x* line mid-season (Fig. 1A). By stem elongation, crops sown early to mid-May received about 0.50 of seasonal rainfall at Hart, Roseworthy and Turretfield, and 0.70 of seasonal rainfall in cooler Mintaro where crops develop more slowly (Figure 1B). The fraction of seasonal rainfall by stem elongation increased to 0.70-0.79 for crops sown early to mid-July. By flowering, crops sown early to mid-May received at least 0.70 of seasonal rainfall and the fraction increased to 0.90 in crops sown early to mid-July (Figure 1B).

**Figure 1.**
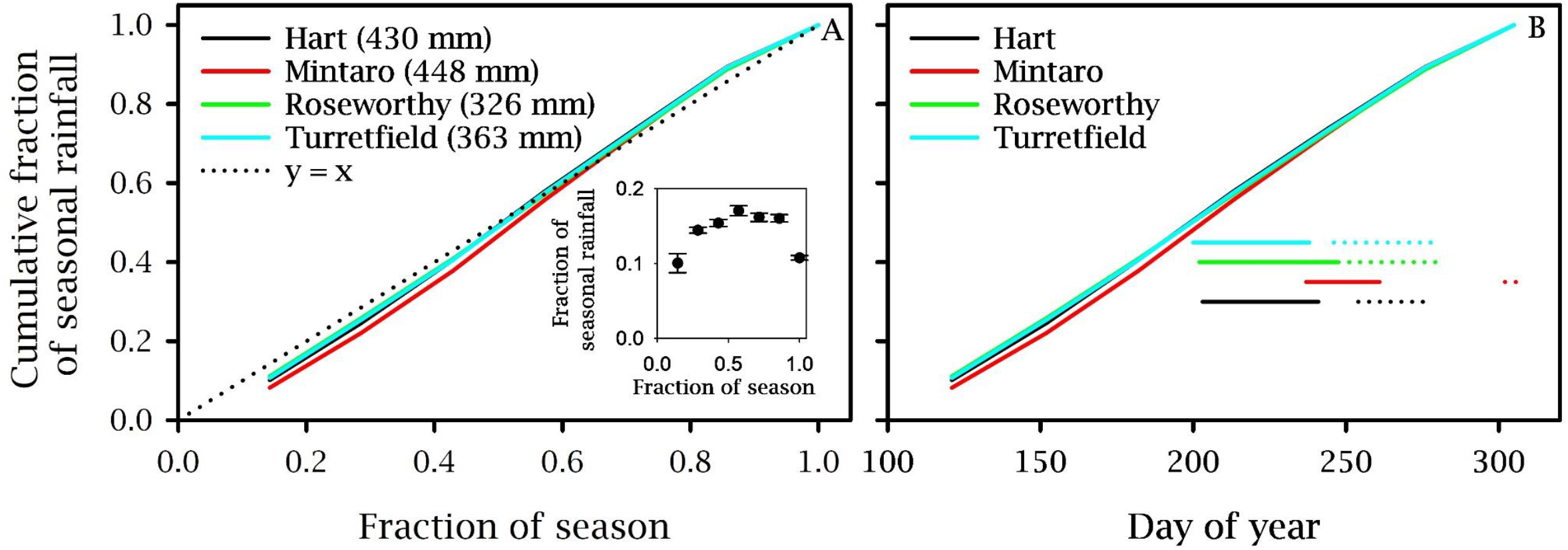
Long-term (1960-2020) cumulative fraction of seasonal rainfall (April to October) in four locations of South Australia as a function of (A) fraction of season and (B) day of year. In (A) average seasonal rainfall is shown between parenthesis and the inset shows the monthly fraction of seasonal rainfall averaged for the four locations, with error bars = 2 standard deviations. In (B) segments are time of stem elongation (solid) and flowering (dashed) where the spread relates to three sources of variation: sowing date (early May to mid-July), variety and their interaction. The late flowering at Mintaro related to frost killing the first cohort of ears, and flowering scored in the second cohort.

### 3.2. Grain yield related to δ^13^C at flowering, booting and stem elongation

Carbon isotope composition measured with IRMS at flowering accounted for 56% of the variation in yield in a sample of 1518 plots with yield from 0.07 to 9.96 t ha^-1^ (Figure 2). Yield also correlated with δ^13^C measured at stem elongation and booting (Figure 3).

**Figure 2.**
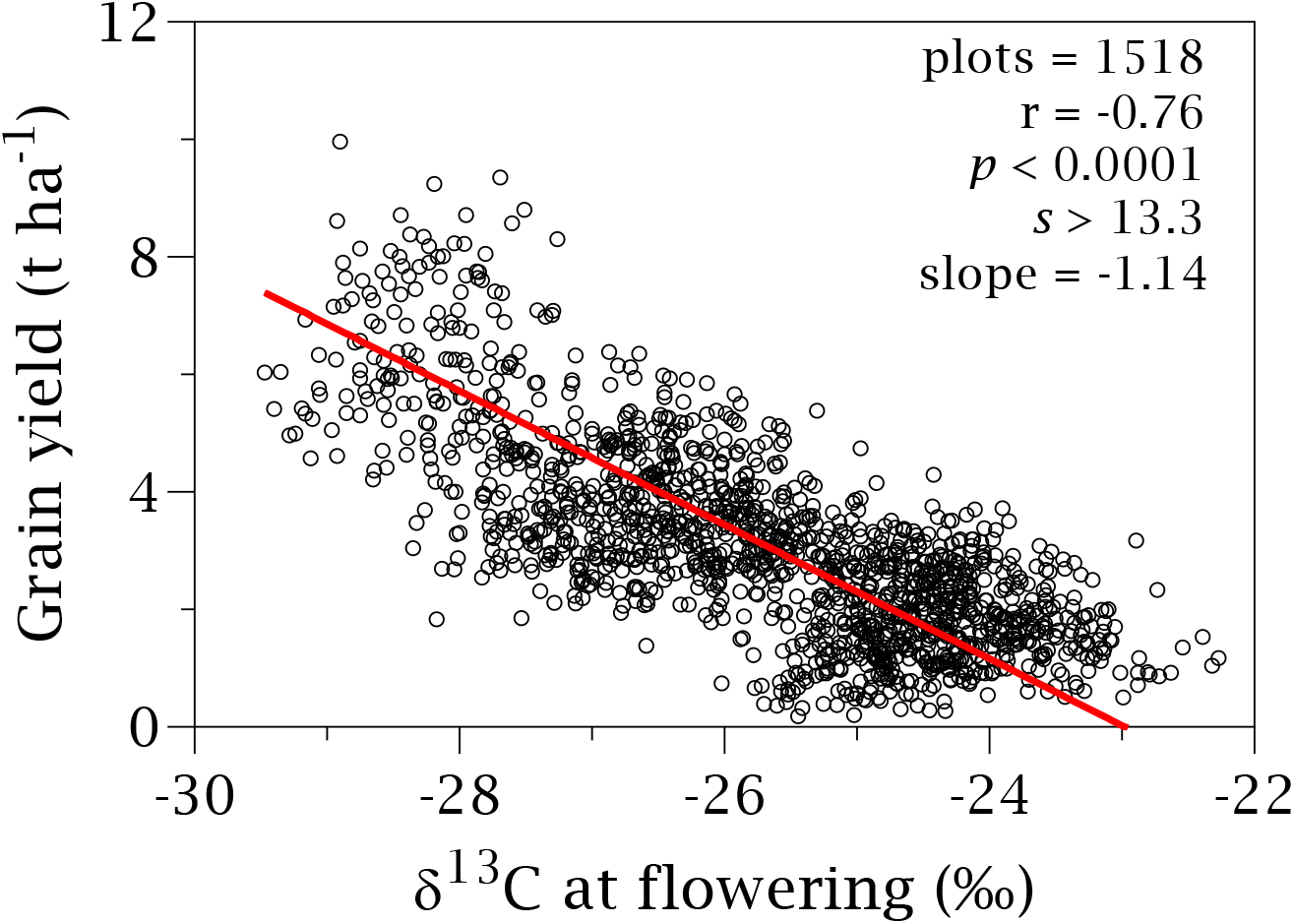
Wheat grain yield correlated with carbon isotope composition (δ^13^C) measured with IRMS at flowering (DC 60 to 68). The line is RMA regression and the slope is in t ha^-1^ ‰^-1^.

**Figure 3.**
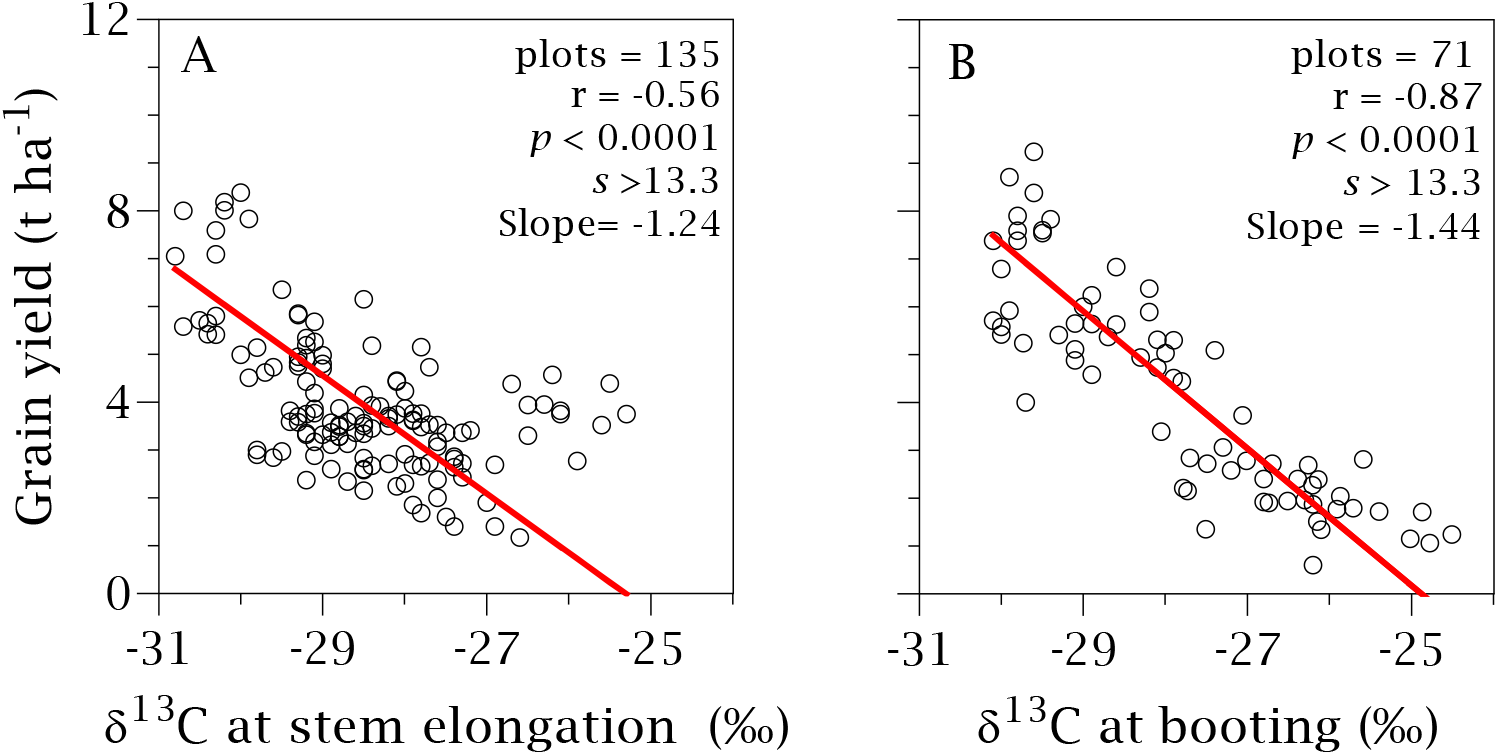
Wheat grain yield correlated with carbon isotope composition (δ^13^C) measured with IRMS at (A) stem elongation (DC 2.3-3.2), and (B) booting (DC 3.7-4.9). Lines are RMA regressions and the slopes are in t ha^-1^ ‰^-1^.

### 3.3. MIR reliably estimated δ^13^C

Global models used a set of 833 samples for calibration and 364 samples for validation comprising three growing stages, 19 cultivars, available nitrogen from 34 to 375 kg ha^-1^, soil available water at sowing from 0 to 161 mm, and growing season rainfall from 118 to 351 mm (Table 1). Figure 4 compares δ^13^C predicted with MIR and δ^13^C measured with IRMS. The calibration PLSR model required 8 factors to explain 82% of the variation in δ^13^C, with RMSEE = 0.48‰. The validation model explained 70% of the variation in δ^13^C, with RMSEP = 0.75‰. The RPD was 2.39‰ for calibration and 1.82‰ for validation.

**Figure 4.**
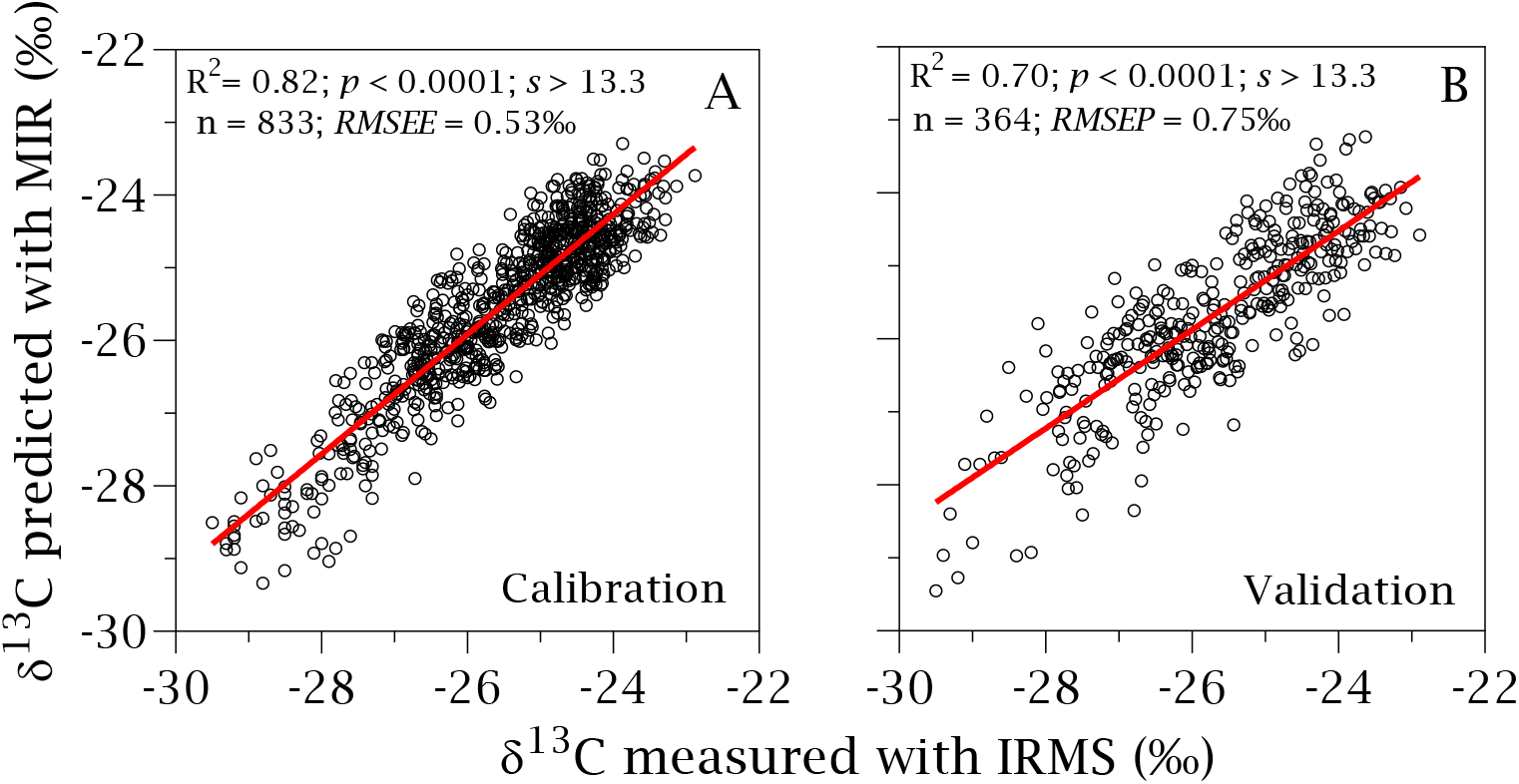
Comparison of δ^13^C predicted with MIR and δ^13^C measured with IRMS in independent (A) calibration and (B) validation sets. Lines are least square regressions.

### 3.4. Yield response to nitrogen associated with δ^13^C at stem elongation in early but not in late-sown crops

The relationship between yield response to nitrogen and predicted δ^13^C with MIR at stem elongation varied with sowing date (Figure 5). For 216 plots sown between 5 and 19 May, yield response to nitrogen varied from -2.0 to 3.5 t ha^-1^ and declined with decreasing *δ*^*13*^C indicative of increasing water stress (Figure 5A). The ranges of both yield and δ^13^C shrank and relationships between yield response to nitrogen and δ^13^C were not apparent with later sowings.

**Figure 5.**
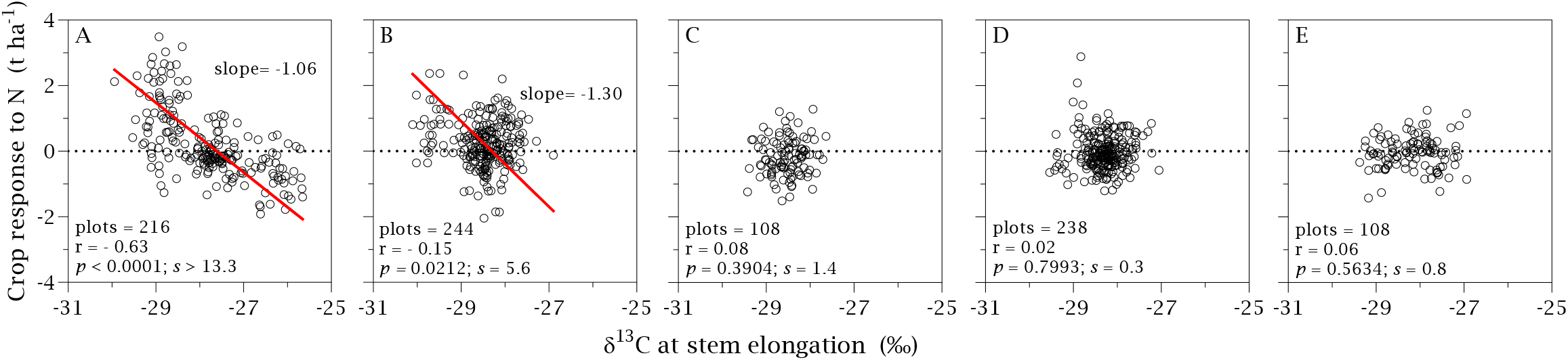
Yield response to nitrogen application as function of δ^13^C predicted with MIR at stem elongation for crops sown at (A) 5-19 May, (B) 20 May-3 June, (C) 4-18 June, (D) 19 June-3 July, (E) 4-20 July. The lines are RMA regressions and the slopes are in t ha^-1^ ‰^-1^.

## 4. Discussion

The physiological interplay between water and nitrogen has agronomic, environmental and economic implications (Sadras et al., 2016). In winter-rainfall environments of south-eastern Australia, the frequency distribution of stored soil water at sowing is L-shaped, with initial soil water contributing a small part of seasonal supply (Sadras and Rodriguez, 2007a; Sadras and Rodriguez, 2007b). For the sampled environments in this study, seasonal rainfall averaged 75% of total water supply (Table 1). Crop reliance on uncertain seasonal rainfall favours a conservative use of nitrogen (Monjardino et al., 2015; Monjardino et al., 2013) causing yield gaps (Hochman and Waldner, 2020; Hochman and Waldner, 2021) and soil mining (Angus and Grace, 2017). In this context, we asked if δ^13^C could contribute to predict yield and yield response to nitrogen, and thus be used to inform fertiliser decisions. The associations between grain yield and δ^13^C is well established in studies with a breeding focus, where genotype and genotype-by-environment interaction are the main sources of variation (Araus et al., 1993; Bort et al., 1998; Condon et al., 2008; Condon and Richards, 1992; Condon et al., 1987; Condon et al., 1992; Ehdaie et al., 1991; Ehdaie and Waines, 1993; Ehdaie and Waines, 1994; Ferrio et al., 2007; Yousfi et al., 2012). Most of these studies focused on δ^13^C measured in grains at maturity, or flag leaves at late stages, rather than in shoot. Here we focus on grain yield and shoot δ^13^C at earlier growing stages where sources of variation are environment and management, with a smaller genotypic component (Table 1). Three findings preliminary support the value of δ^13^C to inform nitrogen management.

First, we found a strong association between yield, in a range from failed crops to 10 t ha^-1^, and δ^13^C at flowering. The physiology of the crop and the prevailing environment explain this association. Yield is a primary function of grain number (Sadras, 2007), and wheat sets grain number in a window from 300 °Cd before to 100 °Cd after anthesis (Dreccer et al., 2007; Fischer, 1985; Telfer et al., 2018). Contrary to the vague notion of terminal drought, the most severe, and one of the more common droughts in Australia, features an onset at ∼400 °Cd before anthesis (about stem elongation) and progresses with the season (Chenu et al., 2013). In our environments, crops received at least 70% of seasonal rainfall by flowering (Figure 1). Hence, δ^13^C at flowering integrates the water status of the crop during the critical period of grain set, and therefore correlates with yield.

Second, we found δ^13^C at stem elongation correlated with yield, and for early-sown crops, with response to nitrogen. The reasons for these associations are less apparent. We can speculate that there is some degree of autocorrelation between water supply early in the season – hence δ^13^C at stem elongation – and seasonal water supply, as suggested in an early analysis of rainfall patterns showing a correlation between April rain and seasonal rain (Sadras et al., 2003).The relationship between yield and δ^13^ C early in the season is less likely where stored soil water at sowing is a larger proportion of seasonal water supply; a field experiment under such conditions showed stable δ^13^ C during the first two months of the growing season, and a gradual deterioration of plant water status in parallel to soil drying and increasing vapour pressure deficit mediated by reduction in stomatal conductance (Condon et al., 1992). Irrespective of the causes, our study preliminary supports the association between yield and δ^13^C at stem elongation necessary to inform nitrogen fertilisation. Furthermore, for a large sample of crops sown in the first half of May, yield response to nitrogen ranged from -2 to 3.5 t ha^-1^, and shifted from positive to negative at δ^13^C = -27.6. The largest response, 3.5 t ha^-1^, was for the crop at Hart with initial soil nitrogen of 111 kg N ha^-1^, initial soil water of 135 mm, 80 mm of rainfall from sowing to stem elongation, and seasonal rainfall of 226 mm (Table 1). Negative yield responses were mostly at Turretfield, with high initial nitrogen (363 kg N ha^-1^), dry soil at sowing (22 mm), 75 mm of rain from sowing to stem elongation, and seasonal rain of 259 mm. In late-sown crops, the range of both δ^13^C and yield response to nitrogen shrank, and relationships disappeared. Consistent with this finding, independent studies in the same region showed a bilinear response of yield to nitrogen supply in May-sown crops, and lack of yield response to nitrogen in late-June crops (Sadras et al., 2019). In late-sown crops, high temperature favouring fast development, short plants and smaller canopies relaxed competition for resources as the driver of grain yield (Sadras et al., 2019).

Third, we showed that MIR spectroscopy can return high-throughput, affordable estimates of δ^13^C of wheat. The error with MIR (RMSEE = 0.53‰, RMSEP = 0.75‰) was higher than the error with IRMS (0.18‰), but δ^13^C estimated with MIR had the resolution required for crop management as demonstrated by correlations with yield response to nitrogen. Previous studies with grains and plant tissues from other species reported δ^13^C estimated with NIR (Ferrio et al., 2001; Kleinebecker et al., 2009). Ferrio et al. (2001) calibrated δ^13^C of durum wheat grain with NIR, reporting an error of 0.46‰ and a coefficient of determination of 0.86, but with a narrower range of environmental variation returning a range of δ^13^C variation that was half of that in our study. Kleinebecker et al. (2009) found coefficients of determination between 0.4 and 0.7 in NIR calibrations of δ^13^C for plant tissues from seven bog species from southern Patagonia, and with variable error depending on the range of variation of δ^13^C for each species. Gouveia et al. (2020) used NIR to estimate the δ^13^C of taro (*Colocasia esculenta* (L.) Schott) and sweet potato (*Ipomoea batatas* (L.) Lam.) tubers and shoots, reinforcing the results of the previous studies using NIR to predict carbon isotopes. Using a single sample, MIR may also be used to estimate other traits such as oxygen and nitrogen isotopes, nutrient deficiency and diseases (Bureau et al., 2019).

## 5. Conclusion

Carbon isotope composition δ^13^C is an agronomically robust trait that correlated with yield not only at flowering but also at stem elongation, the time of nitrogen top dressing. Spectral indices of crop water status are often calibrated against thermal indices, stomatal conductance or leaf water potential. All these traits depend on conditions at the time of measurement (wind, radiation, temperature, vapour pressure deficit) whereas δ^13^C more robustly integrates the water status of the crop from sowing up to the time of measurement. We demonstrate that MIR allows for high throughput, affordable estimation of δ^13^C with the potential to be scaled to paddock level.

## Acknowledgements

We thank GRDC for funding (projects DAS00166; DAS00147); Hart Field Site and the Faulkner family for farm facilities; Barley quality lab, The University of Adelaide, for analytical facilities; Koman Tam (Bruker Pty. Ltd.) for support with MIR; ZH Chow, J Fernández-López, T Lenz, A O’dea, and H Tura for field and lab work; and Australian Grain Technologies for seed.

## CRediT authorship contribution statement

Both authors developed the concept, designed the experiments, analysed data and wrote the paper. Mariano Cossani also carried out part of the field work and all the lab work with isotopes.

## References

Angus, J.F. and Grace, P.R., 2017. Nitrogen balance in Australia and nitrogen use efficiency on Australian farms. Soil Research, 55(6): 435–450.

Anwar, M.R., Rodriguez, D., Liu, D.L., Power, S. and O’Leary, G.J., 2008. Quality and potential utility of ENSO-based forecasts of spring rainfall and wheat yield in south-eastern Australia. Aust. J. Agric. Res., 59(2): 112–126.

Araus, J.L., Amaro, T., Casadesus, J., Asbati, A. and Nachit, M.M., 1998. Relationships between ash content, carbon isotope discrimination and yield in durum wheat. Australian Journal of Plant Physilogy, 25: 835–842.

Araus, J.L., Reynolds, M.P. and Acevedo, E., 1993. Leaf Posture, Grain Yield, Growth, Leaf Structure, and Carbon Isotope Discrimination in Wheat. Crop Science, 33(6): 1273–1279.

Armstrong, R. et al., 2019. Nitrogen supply, rotation and variety are critical predictors of the water use efficiency of wheat in grower’s paddocks in Victoria. In: J. Pratley (Editor), Cells to Satellites. Proceedings of the 19th Australian Society of Agronomy Conference, 25-29 August 2019., Wagga Wagga, NSW, Australia

Asseng, S., McIntosh, P.C., Wang, G. and Khimashia, N., 2012. Optimal N fertiliser management based on a seasonal forecast. European Journal of Agronomy, 38(1): 66–73.

Basso, B., Fiorentino, C., Cammarano, D., Cafiero, G. and Dardanelli, J., 2012. Analysis of rainfall distribution on spatial and temporal patterns of wheat yield in Mediterranean environment. European Journal of Agronomy, 41: 52–65.

Basso, B., Ritchie, J.T., Cammarano, D. and Sartori, L., 2011. A strategic and tactical management approach to select optimal N fertilizer rates for wheat in a spatially variable field. European Journal of Agronomy, 35(4): 215–222.

Bort, J., Araus, J., Hazzam, H., Grando, S. and Ceccarelli, S., 1998. Relationships between early vigour, grain yield, leaf structure and stable isotope composition in field grown barley. Plant Physiology and Biochemistry, 36(12): 889–897.

Bort, J., Belhaj, M., Latiri, K., Kehel, Z. and Araus, J.L., 2013. Comparative performance of the stable isotope signatures of carbon, nitrogen and oxygen in assessing early vigour and grain yield in durum wheat. The Journal of Agricultural Science, 152(3): 408–426.

Bureau, S., Cozzolino, D. and Clark, C.J., 2019. Contributions of Fourier-transform mid infrared (FT-MIR) spectroscopy to the study of fruit and vegetables: A review. Postharvest Biology and Technology, 148: 1–14.

Calderini, D.F. et al., 2021. Overcoming the trade-off between grain weight and number in wheat by the ectopic expression of expansin in developing seeds leads to increased yield potential. New Phytologist, 230(2): 629–640.

Cassman, K.G., Dobermann, A. and Walters, D.T., 2002. Agroecosystems, nitrogen-use efficiency, and nitrogen management. Ambio, 31(2): 132–140.

Chenu, K., Deihimfard, R. and Chapman, S.C., 2013. Large-scale characterization of drought pattern: A continent-wide modelling approach applied to the Australian wheatbelt - spatial and temporal trends. New Phytologist, 198(3): 801–820.

Clark, D.H., Johnson, D.A., Kephart, K.D. and Jackson, N.A., 1995. Near infrared reflectance spectroscopy estimation of 13C discrimination in forages. Society for Range Management, pp. 132-136.

Clark, M. and Tilman, D., 2017. Comparative analysis of environmental impacts of agricultural production systems, agricultural input efficiency, and food choice. Environmental Research Letters, 12(6): 064016.

Condon, A.G. et al., 2008. Stomatal aperture related traits and yield potential in bread wheat. In: M.P. Reynolds, J. Pietragalla and H.-J. Braun (Editors), International symposium of wheat yield potential: Challenges to international wheat breeding. CIMMY, Mexico, DF, pp. 197.

Condon, A.G. and Richards, R.A., 1992. Broad sense heritability and genotype x environment interaction for carbon isotope discrimination in field-grown wheat. Australian Journal of Agricultural Research, 43: 921–934.

Condon, A.G., Richards, R.A. and Farquhar, G.D., 1987. Carbon isotope discrimination is positive correlated with grain yield and dry matter production in field-grown wheat. Crop Science, 27: 996–1001.

Condon, A.G., Richards, R.A. and Farquhar, G.D., 1992. The Effect of Variation in Soil Water Availability, Vapour Pressure Deficit and Nitrogen Nutrition on Carbon Isotope Discrimination in Wheat. Australian Journal of Agricultural Research, 43: 935–947.

Condon, A.G., Richards, R.A. and Farquhar, G.D., 1993. Relationships between carbon isotope discrimination, water use efficiency and transpiration efficiency for dryland wheat. Aust. J. Agric. Res., 44: 1693–1711.

Cossani, C.M. and Sadras, V.O., 2021. Nitrogen and water supply modulate the effect of elevated temperature on wheat yield. European Journal of Agronomy, 124.

Dreccer, M.F., Borgognone, M.G., Ogbonnaya, F.C., Trethowan, R.M. and Winter, B., 2007. CIMMYT-selected derived synthetic bread wheats for rainfed environments: Yield evaluation in Mexico and Australia. Field Crops Research, 100(2-3): 218–228.

Ehdaie, B., Hall, A.E., Farquhar, G.D., Nguyen, H.T. and Waines, J.G., 1991. Water-use efficiency and carbon isotope discrimination in wheat. Crop Science, 31: 1282–1288.

Ehdaie, B. and Waines, J.G., 1993. Variation in Water-Use Efficiency and Its Components in Wheat: I. Well-Watered Pot Experiment. Crop Science, 33(2): cropsci1993.0011183X003300020016x.

Ehdaie, B. and Waines, J.G., 1994. GROWTH AND TRANSPIRATION EFFICIENCY OF NEAR-ISOGENIC LINES FOR HEIGHT IN A SPRING WHEAT. Crop Science, 34(6): 1443–1451.

Farquhar, G.D. and Richards, R.A., 1984. Isotopic composition of plant carbon correlates with water-use efficiency of wheat genotypes. Australian Journal of Plant Physiology, 11(6): 539–552.

Ferrio, J.P., Bertran, E., Nachit, M., Royo, C. and Araus, J.L., 2001. Near infrared reflectance spectroscopy as a potential surrogate method for the analysis of Δ^13^C in mature kernels of durum wheat. Australian Journal of Agricultural Research, 52(8): 809–816.

Ferrio, J.P. et al., 2007. Relationships of grain δ13C and δ18O with wheat phenology and yield under water-limited conditions. Annals of Applied Biology, 150(2): 207–215.

Fischer, R.A., 1985. Number of kernels in wheat crops and the influence of solar radiation and temperature. The Journal of Agricultural Science, 105(2): 447–461.

French, R.J. and Schultz, J.E., 1984. Water use efficiency of wheat in a Mediterranean type environment. I. The relation between yield, water use and climate. Australian Journal of Agricultural Research, 35: 743–764.

Gentilli, J., 1971. The main climatological elements. In: J. Gentilli (Editor), Climates of Australia and New Zealand. World Survey of Climatology. Elsevier, New York, pp. 119–188.

Gouveia, C.S.S., Lebot, V. and Pinheiro de Carvalho, M., 2020. NIRS Estimation of Drought Stress on Chemical Quality Constituents of Taro (Colocasia esculenta L.) and Sweet Potato (Ipomoea batatas L.) Flours.,. Appl. Sci., 10: 8724.

Greenland, S., 2019. Valid P-Values Behave Exactly as They Should: Some Misleading Criticisms of P-Values and Their Resolution With S-Values. The American Statistician, 73(sup1): 106–114.

Hall, A.E., Richards, R.A., Condon, A.G., Wright, G.C. and Farquhar, G.D., 1994. Carbon isotope discrimination and plant breeding. Plant Breed. Rev., 12: 81–113.

Hochman, Z. and Waldner, F., 2020. Simplicity on the far side of complexity: optimizing nitrogen for wheat in increasingly variable rainfall environments. Environmental Research Letters, 15(11): 114060.

Hochman, Z. and Waldner, F., 2021. Corrigendum: Simplicity on the far side of complexity: optimizing nitrogen for wheat in increasingly variable rainfall environments (2020 Environ. Res. Lett. 15 114060). Environmental Research Letters, 16(4): 049501.

Hoogmoed, M., Neuhaus, A., Noack, S. and Sadras, V., 2018. Benchmarking wheat yield against crop nitrogen status, 222, 153–163 pp.

Hoogmoed, M. and Sadras, V.O., 2018. Water Stress Scatters Nitrogen Dilution Curves in Wheat. Frontiers in Plant Science, 9: 406.

Jobbágy, E.G. and Sala, O.E., 2014. The imprint of crop choice on global nutrient needs. Environmental Research Letters, 9(8).

Kleinebecker, T., Schmidt, S.R., Fritz, C., Smolders, A.J.P. and Hölzel, N., 2009. Prediction of δ13C and δ15N in plant tissues with near-infrared reflectance spectroscopy. New Phytologist, 184(3): 732–739.

Kohn, M.J., 2010. Carbon isotope compositions of terrestrial C3 plants as indicators of (paleo)ecology and (paleo)climate. Proceedings of the National Academy of Sciences of the United States of America, 107(46): 19691–19695.

Ludbrook, J., 2012. A primer for biomedical scientists on how to execute Model II linear regression analysis. Clinical and Experimental Pharmacology and Physiology, 39(4): 329–335.

Meier, E.A., Hunt, J.R. and Hochman, Z., 2021. Evaluation of nitrogen bank, a soil nitrogen management strategy for sustainably closing wheat yield gaps. Field Crops Research, 261: 108017.

Moeller, C., Smith, I., Asseng, S., Ludwig, F. and Telcik, N., 2008. The potential value of seasonal forecasts of rainfall categories - Case studies from the wheatbelt in Western Australia’s Mediterranean region. Agricultural and Forest Meteorology, 148(4): 606–618.

Monjardino, M., McBeath, T., Ouzman, J., Llewellyn, R. and Jones, B., 2015. Farmer risk-aversion limits closure of yield and profit gaps: A study of nitrogen management in the southern Australian wheatbelt. Agricultural Systems, 137: 108–118.

Monjardino, M., McBeath, T.M., Brennan, L. and Llewellyn, R.S., 2013. Are farmers in low-rainfall cropping regions under-fertilising with nitrogen? A risk analysis. Agricultural Systems, 116: 37–51.

Neuhaus, A. and Sadras, V.O., 2018. Relationship between rainfall-adjusted nitrogen nutrition index and yield of wheat in Western Australia. Journal of Plant Nutrition ():.

Okano, K., Ito, O., Kokubun, N. and Totsuka, T., 1983. Determination of 13C in plant materials by infrared absorption spectrometry using a simple calibration method. Soil Science and Plant Nutrition, 29(3): 369–374.

Pancorbo, J.L. et al., 2021. Simultaneous assessment of nitrogen and water status in winter wheat using hyperspectral and thermal sensors. Eur. J. Agron., 127: 126287.

Rebetzke, G.J., Condon, A.G., Richards, R.A. and Farquhar, G.D., 2002. Selection for reduced carbon isotope discrimination increases aeral biomass and grain yield of rainfed bread wheat. Crop Science, 42: 739–745.

Richards, R.A., López-Castañeda, C., Gomez-Macpherson, H. and Condon, A.G., 1993. Improving the efficiency of water use by plant breeding and molecular biology. Irrigation Science, 14(2): 93–104.

Sadras, V., Roget, D. and Krause, M., 2003. Dynamic cropping strategies for risk management in dry-land farming systems. Agricultural Systems, 76(3): 929–948.

Sadras, V.O., 2007. Evolutionary aspects of the trade-off between seed size and number in crops. Field Crops Research, 100(2-3): 125–138.

Sadras, V.O. et al., 2016. Interactions between water and nitrogen in Australian cropping systems: physiological, agronomic, economic, breeding and modelling perspectives. Crop and Pasture Science, 67: 1019–1053.

Sadras, V.O. and Rodriguez, D., 2007a. The limit to wheat water-use efficiency in eastern Australia. II. Influence of rainfall patterns. Australian Journal of Agricultural Research, 58(7): 657–669.

Sadras, V.O. and Rodriguez, D., 2007b. Seasonality can be the difference between wet and dry. Ground Cover, 68: 29.

Sadras, V.O., Thomas, D., Cozzolino, D. and Cossani, C.M., 2019. Wheat yield response to nitrogen from the perspective of intraspecific competition. Field Crops Research, 243.

Schulze, E.-D., Turner, N.C., Nicolle, D. and Schumacher, J., 2006a. Species differences in carbon isotope ratios, specific leaf area and nitrogen concentrations in leaves of Eucalyptus growing in a common garden compared with along an aridity gradient. Physiologia Plantarum, 127(3): 434–444.

Schulze, E.D., Turner, N.C., Nicolle, D. and Schumacher, J., 2006b. Leaf and wood carbon isotope ratios, specific leaf areas and wood growth of Eucalyptus species across a rainfall gradient in Australia. Tree Physiology, 26(4): 479–492.

Slafer, G.A., Savin, R. and Sadras, V.O., 2014. Coarse and fine regulation of wheat yield components in response to genotype and environment. Field Crops Research, 157: 71–83.

Telfer, P. et al., 2018. A field and controlled environment evaluation of wheat (Triticum aestivum) adaptation to heat stress. Field Crops Research, 229: 55–65.

Yousfi, S., Serret, M.D., Marquez, A.J., Voltas, J. and Araus, J.L., 2012. Combined use of delta(1)(3)C, delta18O and delta15N tracks nitrogen metabolism and genotypic adaptation of durum wheat to salinity and water deficit. New Phytol, 194(1): 230–44.

Zadoks, J.C., Chang, T.T. and Konzak, C.F., 1974. A decimal code for the growth stages of cereals. Weed Res., 14: 415–421.

